# Assessment of autophagy in *Leishmania* parasites

**DOI:** 10.1101/2024.01.03.574013

**Authors:** Somtochukwu S. Onwah, Jude E. Uzonna, Saeid Ghavami

**Affiliations:** Department of Immunology, Max Rady College of Medicine, University of Manitoba, Winnipeg, MB R3E 0T5 Canada; Department of Medical Microbiology and Infectious Diseases, Max Rady College of Medicine, University of Manitoba, Winnipeg, MB R3E 0T5 Canada; Department of Pathology, Max Rady College of Medicine, University of Manitoba, Winnipeg, MB R3E 0T5 Canada; Biology of Breathing Theme, Children Hospital Research Institute of Manitoba, Winnipeg, MB R3E 0V9, Canada; University of Manitoba, Winnipeg, MB R3E 0V9, Canada; Biology of Breathing Theme, Children Hospital Research Institute of Manitoba, University of Manitoba, Winnipeg, MB R3E 0V9, Canada; Academy of Silesia, Faculty of Medicine, Rolna 43, 40-555 Katowice, Poland; Research Institutes of Oncology and Hematology, Cancer Care Manitoba-University of Manitoba, Winnipeg, MB R3E 0V9, Canada; Department of Human Anatomy and Cell Science, University of Manitoba College of Medicine, Winnipeg, MB R3E 0V9, Canada

**Author notes:** Correspondence: Dr. Saeid Ghavami, Somtochukwu S. Onwah.

**Keywords:** Cutaneous leishmaniasis, *Leishmania major*, Autophagy, TEM, Acridine Orange

## Abstract

Leishmaniasis is a neglected tropical disease caused by numerous species of *Leishmania* parasites, including *Leishmania major.* The parasite is transmitted by several species of sandfly vectors and infects myeloid cells leading to a myriad of inflammatory responses, immune dysregulations, and disease manifestations. Every cell undergoes autophagy, a self-regulated degradative process that permits the cells to recycle damaged or worn-out organelles in order to maintain cellular health and homeostasis. Studies have shown that *Leishmania* modulates their host cell autophagic machinery and there are indications that the parasite-specific autophagic processes may be valuable for parasite virulence and survival. However, the role of autophagy in *Leishmania* is inconclusive because of the limited tools available to study the *Leishmania*-specific autophagic machinery. Here, we describe methods to study and definitively confirm autophagy in *Leishmania major*. Transmission electron microscopy (TEM) allowed us to visualize *Leishmania* autophagosomes, especially those containing damaged mitochondrial content, as well as dividing mitochondria with ongoing fusion/fission processes. Flow cytometry enabled us to identify the amount of acridine orange dye accumulating in the acidic vacuolar compartments in *Leishmania major* by detecting fluorescence in the red laser when autophagic inhibitors or enhancers were included. These methods will advance studies that aim to understand autophagic regulation in *Leishmania* parasites that could provide insights into developing improved therapeutic targets against leishmaniasis.

## 1 Introduction

Autophagy is a natural “in-house” cleaning process that allows a stressed cell to breakdown cellular components and recycle/reuse some parts for efficient cellular processes (1, 2). Many cells require autophagy to recycle damaged or stressed intracellular components for the purpose of maintaining homeostasis and cell health through both selective or non-selective degradative mechanisms (3, 4). The autophagic process is activated when the cell senses a need for nutrients, growth or energy (5–7) or to eliminate invasive pathogens (8–10). All cells are known to engage in basal levels of autophagy to perform basic hemostatic processes that include protein or organelle turnover (8, 11). In mammals, the induction of autophagy is a complicated process that involves a myriad of autophagy-like proteins, ATGs, which induce the formation of the autophagic organelle, autophagosome. Once the autophagosomes mature, they engulf many intracellular materials, transporting them to the lysosome for further degradation (12). Autophagy evolved to help eliminate invading intracellular pathogens by releasing reactive oxygen species and degradative proteolytic enzymes. It directs the imbalance in the nutrient availabilities and cations, which causes the destruction of organisms (13). In line with this, the continuous persistence of intracellular pathogens is associated with manipulating the host autophagic pathway (14, 15). Indeed, the establishment of many infections and human disease pathologies have been linked to an aberrant autophagic machinery (16–18). Thus, a good understanding of the autophagic processes and targeting the host autophagic machinery could lead to the development of novel therapeutic strategies for controlling disease progression.

Leishmaniasis is a neglected disease that has become a global health burden, with an estimated 1 million new cases occurring annually and over 1 billion individuals living in the endemic continents (such as Africa, Asia, the Americas and the Middle East) at risk (19–21). The disease, which is caused by several species of the protozoan parasite belonging to the genus *Leishmania*, displays diverse clinical manifestations. Transmission of the parasite to their mammalian hosts is through the bite of an infected sandfly vector (19, 21). Cutaneous leishmaniasis is the more common and mild form of the disease and presents as a self-resolving lesion, often resulting in complicated and disfiguring outcomes. The more fatal and severe form of the disease, visceral leishmaniasis, is caused by *L. donovani* and systemically affects multiple organs and if left untreated can lead to death (19, 21). Besides the clinical manifestations, *Leishmania* infections result in chronic inflammation, immune cell dysregulation and cytokine imbalances (22). Despite the myriad of research aimed at understanding interactions between the host cells and *Leishmania* parasites, there is still no approved vaccine against the disease because this complex interaction is poorly understood. Therefore, drug treatment remains the only viable option to control the disease, but the current treatment modalities are plagued with various challenges (23, 24). A clear understanding of the parasite’s cellular biologic processes and molecules that influence and/or drive protective responses against the host could advance the development of novel therapeutic strategies against leishmaniasis (25, 26).

In addition to evading host immune responses, *Leishmania* parasites have been shown to hijack the host cell’s autophagic machinery (8). It has been suggested that the continued survival of *Leishmania* parasites in their host is due to their ability to modulate host mTOR-regulated autophagy processes (27), which helps them to obtain essential nutrients in their harsh intracellular host environment (28). Interestingly, as their mammalian host cells, *Leishmania* parasites have been reported to undergo autophagic processes and nutrient deprivation was observed to enhance autophagy in *Leishmania* (29, 30). Furthermore, the transition of promastigotes (insect) to amastigotes (mammalian) life forms of the parasite depends on autophagy (31), suggesting that just like mammalian cells, autophagic processes are indeed critical for parasite homeostasis. Interestingly, the expression of autophagy-like proteins observed in mammalian cells has also been identified in *Leishmania* parasites (32). ATG4 and ATG8 are some of the well-characterized autophagic proteins that have been identified to play a role in the survival and virulence of *Leishmania* parasites (33–36). The presence of these key autophagic proteins in *Leishmania* suggests ample avenues to target either the host or parasite autophagic machinery for anti-leishmanial interventions. However, not many studies have explored *Leishmania*’s autophagic responses extensively as the exact mechanisms of autophagy in *Leishmania* remains elusive. Therefore, to help understand the mechanisms of autophagy in *Leishmania*, tools geared at measuring autophagy in the parasites are needed.

Here, we describe two complementary methods for detecting autophagy in *Leishmania*. Transmission electron microscopy (TEM) has long been the tool that accurately detects autophagic accumulation and organelles in mammalian cells because the exact ultrastructural composition is revealed in nanoscales (37). This paper reports the visual identification of specific autophagic organelles and structures in *Leishmania major* by TEM. During autophagy, there is an increase in the formation of acidic vacuolar organelles (AVO), such as lysosomes and auto phagolysosomes (38). Acridine orange (AO) is known to accumulate in these acidic compartments and fluoresces bright red (38). However, since the use of multiple methods to detect autophagy is encouraged, (39) we used flow cytometry to detect and validate the accumulation of AO in the AVO of *Leishmania major*. Although TEM is the gold standard for accessing autophagy because of the high-resolution images and morphometric output (37, 39, 40), in assessing autophagic flux, a time-lapse observation is critical to deduce autophagic mechanisms which is not achievable with TEM (40). Live cell imaging through fluorescence microscopy is an alternative tool that measures autophagic flux (41). Also, identification of autophagic proteins such as the accumulation of MAP1LC3/LC3-II in western blots is considered an indication of autophagosomal flux especially in the presence of inhibitors of the autophagic process such as bafilomycin A1 (42). Assessing autophagic flux in *Leishmania* has been challenging because there are no anti-autophagic protein antibodies such as anti-LC3 or anti-p62. Thus, accessing autophagic flux by measuring the accumulation of acridine orange in the AVO of *Leishmania* over time is a technique that could be critical to unravel the mechanisms of autophagy in *Leishmania*.

## 2 Materials

Aseptic techniques were applied in the preparation of *Leishmania* parasites in axenic culture, and all work with the parasites was carried out inside a Biosafety hood in a Biosafety level 2 laboratory.

### 2.1 *Leishmania* parasites growth and media

1. *Leishmania major* MHOM/80/Friedlin strain
2. Twenty Percent (20%) Inactivated-Fetal Bovine Serum (FBS)
3. Medium 199 (M199) incomplete; mix 9.5 g M199 powder (Gibco, Cat# 31100035), 2.2g NaHCO_3_ (Sigma Cat# S6014-25G), 25 ml of 1 M Hepes (Gibco, Cat# 15630080), 10 mM Adenine in 50 mM Hepes, 0.25% Hemin in 50% Triethanolamine, 0.1% Biotin in 95% ethanol, 0.25 mg/ml Biopterin in PBS, and distilled water up to a total volume of 1L
4. T25 Biolite™ culture flasks (ThermoFischer, Cat# 130189)
5. Sterile pipette tips and pipettors
6. Incubator set at 27 °C
7. Penicillin-Streptomycin (Gibo™ Cat # 15140122)
8. Water bath set at 37 °C
9. Centrifuge 4000 rpm speed max (ThermoFischer Cat# 75007200)
10. Optical Microscope
11. Glass vacuum filtration apparatus
12. Fifteen millilitres (15 ml) Falcon tubes
13. 1L and 200 ml sterile bottles

### 2.2 Fixing parasites for Electron Microscopy

1. 6% Glutaraldehyde stock stored at 4 °C (see Note 1)
2. 0.2 M Sorensen’s Phosphate buffer stored at 4 °C
3. 0.5 M Sorensen’s Phosphate buffer store at room temperature (RT) and should be a clear solution.
4. Sucrose powder
5. Fixation solution; 0.1 M Sorensen’s phosphate buffer with 3% Glutaraldehyde (prepare a 1:1 mixture of 0.2 M Sorensen’s buffer and 6% glutaraldehyde (see Note 2 & 3)
6. Sucrose solution: prepare 5% Sucrose in 0.1 Sorensen’s phosphate buffer for instance a final volume of 25 ml, mix 1.25g of sucrose with 5ml of 0.5M Sorensen’s Phosphate buffer up to 25 ml of ddH_2_O (see Note 2).
7. Centrifuge (Fisher Scientific Cat# 13864456)
8. Fifteen (15), 1.5 ml Eppendorf tubes
9. Philips-CMC-300 KV Transmission Electronic Microscope (Netherlands), Histology lab, University of Manitoba, Canada.
10. 1X Phosphate buffered saline (PBS)

### 2.4 Experiments with Acridine orange

1. Acridine orange hydrochloride solution (Sigma, Cat# A8097)
2. ddH_2_O
3. 5 ml conical glass tubes
4. Tube racks
5. Foil paper.
6. 100 nM Bafilomycin
7. 2% FBS
8. 1X Phosphate buffered saline (PBS)
9. 24 well plate

### 2.3 Flow cytometry

1. Beckman Coulter CytoFlex LX Digital Flow Cytometry Analyzer Four Laser System at the University of Manitoba, Flow cytometry Core facility.
2. Analysis on Flow jo Version 10 software (TreeStar, Ashland, OR)

## 3 Methods

### 3.1 Preparing parasite complete media

1. M199 incomplete media is prepared and filtered using the vacuum filtration system inside a biosafety hood. After filtration, the media is poured into a 1L sterile bottle.
2. For 100 ml complete M199 media, add 80 ml of the incomplete M199 media in a 200 ml sterile bottle alongside 20 ml of FBS (heat-inactivated) and 1 ml of 1000 U/ml Penicillin-Streptomycin to prevent bacteria contamination.
3. Shake gently by swirling the bottle in a circular motion to mix.

### 3.2 Preparing *Leishmania* axenic cultures

1. *L. major* parasites MHOM/80/Friedlin strain stored in cryo vials in Liquid Nitrogen are swirled in 37 °C water bath until thawed (see Note 4)
2. 10 ml of complete M199 media is poured into a 15 ml Falcon tube and the thawed contents of the cryo vial from Step 1 above is added.
3. Centrifuge at 3000 rpm for 15 minutes
4. Label T25 tubes and add fresh 10 ml complete M199 medium
5. Discard supernatant from step 3, pipette out the pellets and add them into the T25 flash containing complete M199 medium.
6. Visually check to see parasites in the Flask using an optical microscope (Note 5)
7. Incubate at 27 °C for 3 days
8. Under an optical microscope, check that the parasites are healthy, motile and have their characteristic slender body with a flagellum.
9. Passage the parasites by taking 100 μl into a new T25 flask containing 10 ml of complete M199 medium and further incubate at 27 °C. At the specified days post-passage, the parasites are ready for downstream experiments.

### 3.3 Fixation of parasites for Transmission Electron Microscopy imaging

1. At day 3 and 7 post-passage, the parasites were transferred into a 15 ml Falcon tube inside a biosafety cabinet.
2. Centrifuge at 3000 rpm for 7 minutes (see Note 6).
3. Discard supernatant and transfer pellets containing the parasites into 1.5 ml Eppendorf tubes.
4. Resuspend the parasites in PBS and gently stir by pipetting up and down. Do not vortex.
5. Spin at 3000 rpm for 7 minutes.
6. Discard the supernatant and add 1 ml of the Fixation solution. Let the parasites be fully resuspended for proper fixation by gently pipetting up and down.
7. Fix the parasites for 3 hours inside a fume hood at RT
8. Pellet the parasites by spinning at 3000 rpm for 7 minutes, remove as much of the fixative as possible and replace them with 1 ml of 5% sucrose in 0.1 Sorensen’s buffer. Keep parasites at 4 °C
9. Fixed parasites are embedded unto the stage of an electron microscope following the instructions of the transmission electron microscopy manual. A trained histologist should identify parasite containing double membrane vacuoles, mitochondria in the parasite, ER in the parasite, and the cytosolic membrane after capturing images at 34000x magnification.

### 3.4 Acridine orange stain and Flow cytometry acquisition

1. At day 3, passaged parasites were counted using a hemocytometer or a cell counter.
2. In some wells, plate 1 x 10^6^ parasites in 1 ml complete M199 medium and in other wells plate 1 x 10^6^ parasites containing incomplete M199 medium supplemented with 100 U/ml penicillin, 100 μg/ml streptomycin and 1% FBS in a 24-well plate (See Note 7).
3. After 48 hrs post-day 3 and day 7 cultures, 100 nM Bafilomycin was added to some of the wells that contained complete M199 medium.
4. At the time point to measure autophagy (3 and 7 days) parasites in individual wells of the 24 well plate were transferred to a 5 ml glass conical tube.
5. Wash parasites twice in PBS by spinning at 3000 rpm for 10 minutes.
6. Discard supernatants and add 1 μl of 1 mg/ml acridine orange solution to the pellets in the dark at RT for 10 minutes (See Note 8).
7. Wash parasites twice in PBS by spinning at 3000 rpm for 10 minutes and resuspended in 200 μl of PBS for acquisition in the cytoflex machine
8. Acquisition was done by opening channels in the red (PECy5) and green (FITC) lasers and recording 20000 events.
9. Further analysis of acridine orange accumulating in the lysosomal vesicles was assessed by the ratio of the mean florescence intensity (MFI) in the PECy5 channel to the FITC channel by the flowjo software.

### 3.5 Results

#### 3.5.1 Visualizing autophagic organelles by TEM

The TEM images of fixed *L. major* captured double membranous autophagosomes including those containing damaged mitochondrial contents, as well as mitochondria with an ongoing fusion/fission process (**Figure 1**). These images suggest that *Leishmania* undergoes basal autophagic processes (quality control processes) that enable the removal of damaged mitochondria.

**Figure 1:**
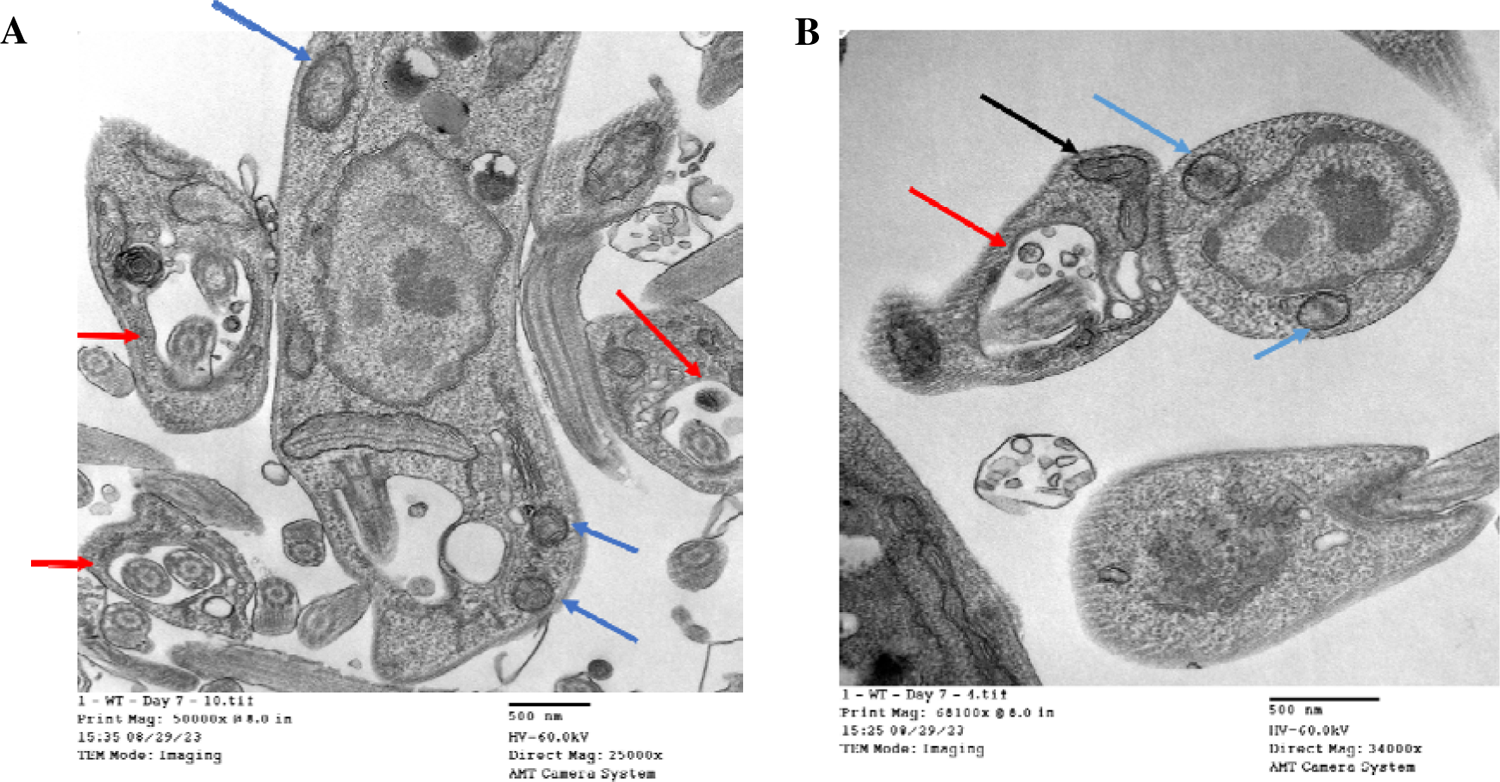
Images of autophagic organelles in Leishmania major by TEM. Fixed Leishmania major at day 7 of axenic culture growth showing double membrane vacuoles representing autophagosome, (A&B, Blue arrows), autophagosome containing damaged mitochondrial (A&B,, red arrows) and dividing mitochondria with an ongoing fusion/fission process (B, black arrows). Images were taken with a magnification of 34000X.

#### 3.5.2 Accumulation of acridine orange in the lysosomal vacuoles by flow cytometry

The strategy employed in the acquisition of the AO-stained parasites by flow cytometry was to gate on live parasite from the forward and live scatter axis plot. Following gating on single parasites (singlets), the red and green florescence in the PECy5 and FITC channels were gated, respectively (Figure 2A). The frequency and ratios of the mean fluorescent intensity of the AO stained parasites in the red and green florescent channels without treatment, with 100 nM Bafilomycin (autophagy inhibition) and in nutrient starvation condition (1% FBS, autophagy enhancer), were assessed (Figure 2B**& C**). In the parasites, basal autophagic processes were occurring in the no treatment group because when autophagy is inhibited by Bafilomycin, there was a decrease in the mean florescent intensity (MFI) of parasites gated in the red channel. However, in the nutrient deprivation condition (1% FBS) which enhances autophagy, there was a significant increase in the MFI of the red florescence compared to the untreated parasite groups (Figure 2B**& C**).

**Figure 2:**
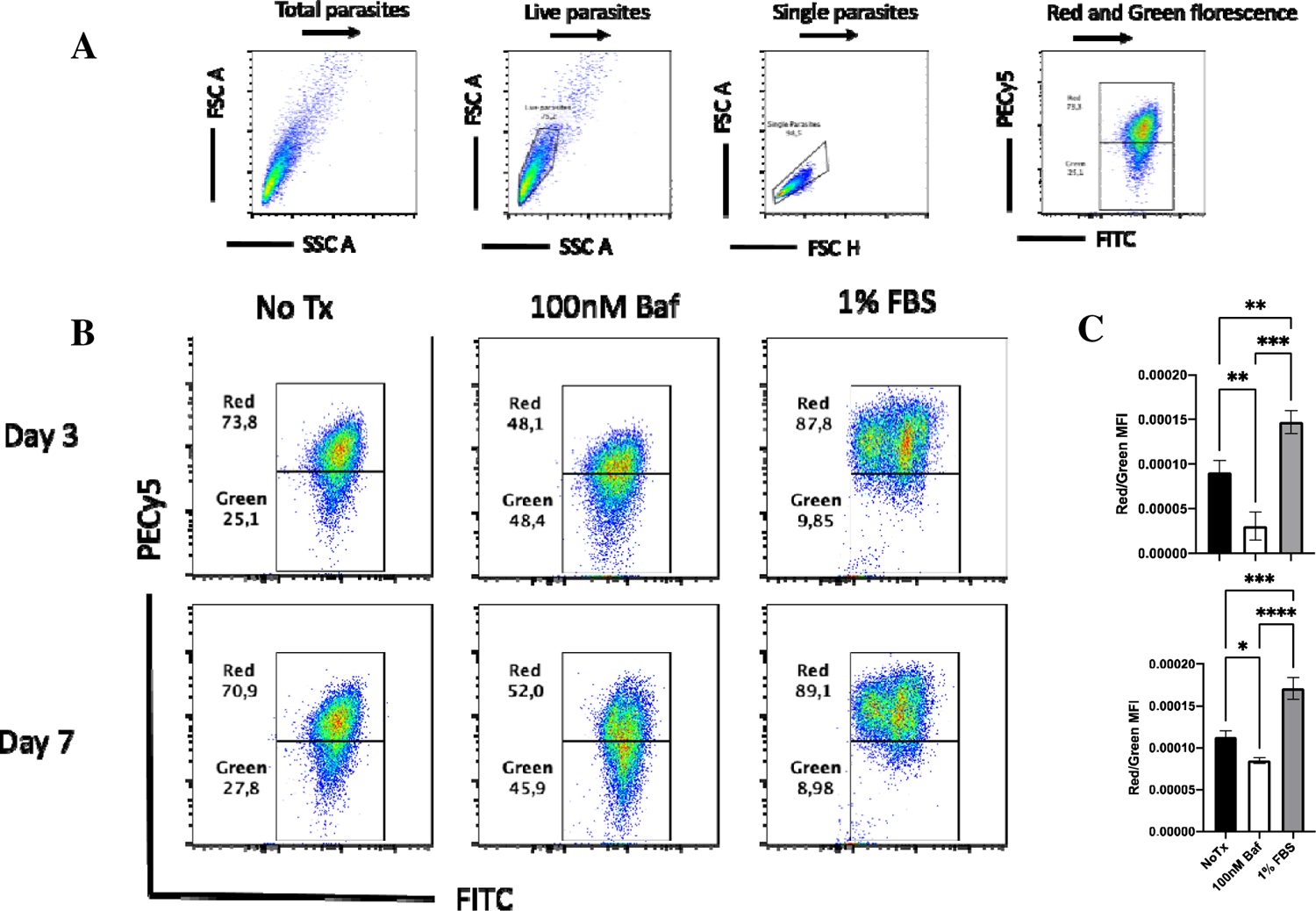
Acridine orange accumulating in the lysosomal vacuole in Leishmania major by Flow cytometry. Axenic cultures of Leishmania major at day 3 and 7 were stained with acridine orange and the amount of red florescence detected by flow cytometry indicates autophagic activity occurring in the lysosomes. (A) Gating strategy (B &C) flow plots and graphs demonstrating the ratio of the MFI of the red florescence to green florescence when the parasites have no treatment, given autophagic inhibitors (Bafilomycin, Baf), or exposed to a nutrient deprivation model (1% FBS).

#### Notes

1. Use if it is completely clear but once it starts turning yellow, discard and make a fresh batch.
2. Prepare fresh on the day of fixation
3. Work in the fume hood
4. All parasite media preparation and cultures should be carried out in a Biosaftety hood.
5. Morphologically, the parasites may have an oval shape with less pronounced flagella when brought straight out from liquid nitrogen. Occasional movements may be observed with these parasites at this time.
6. Too high spins and longer periods can damage the parasite’s ultrastructure.
7. This was carried out for two time points day 3 and day 7 parasite cultures in separate plates to prevent the risk of contamination as the experiments were done on different days.
8. The tubes were placed in a rack and covered with foil paper but alternatively can be kept in a dark cupboard because acridine orange is light sensitive and can easily degrade.

## Acknowledgement

SG conceptualized and designed the study. SSO performed the experiments. SG and SSO analyzed the data. SSO wrote the manuscript. SSO, SG, JU reviewed and edited the manuscript. JU funded and SG supervised the experiments.

